# A premotor microcircuit to generate behavior-specific muscle activation patterns in Drosophila larvae

**DOI:** 10.1101/2022.08.18.504452

**Authors:** Yuhan Huang, Aref A Zarin

**Affiliations:** Department of Biology, Texas A&M University, College Station, TX

**Keywords:** larval locomotion, multifunctional motor circuits, connectome, motor neuron, premotor neuron, Drosophila, muscle calcium imaging, ssTEM

## Abstract

Animals can use a common set of muscles and motor neurons (MNs) to generate diverse locomotor behaviors, but how this is accomplished remains poorly understood. Previously, we characterized the muscle activity patterns for Drosophila larval forward and backward locomotion and found that ventral oblique (VO) muscles become active earlier in backward than in forward locomotion (Zarin et al. 2019). Here, we describe how premotor circuits generate differential activation timing of VO muscles. We identify inhibitory (A06c) and excitatory (A27h) premotor neurons (PMNs) with the greatest number of synapses with VO MNs. Strikingly, A06c is a bi-modal PMN that fires before and after VO MNs in forward locomotion but fires only after MNs in backward locomotion. Further, A27h is a forward-dedicated PMN active only in forward locomotion. These two PMNs interconnect with another forward-dedicated excitatory PMN (A18b3), to create feedforward inhibitory microcircuits that define the activity window for VO MNs/muscles, producing precise VO muscle patterns underlying forward locomotion. Silencing A06c, A27h, or A18b3 PMN results in premature VO muscle activation in forward locomotion, resembling early VO activation in backward locomotion. Our results identify PMN micro-circuits that produce unique MN/muscle activity patterns to create behavior-specific motor output.

## Introduction

Motor circuits are multifunctional and can generate different motor outputs, enabling the animals perform diverse locomotor behaviors using the same muscles. The mechanism underlying the multifunctionality of motor circuits remains a fundamental yet open question.

A possible mechanism for generation of such distinct motor outputs is that the same MN may receive behavior-specific inputs from its upstream premotor interneurons (PMNs). This theory is supported by discovery of interneurons that are dedicated to a specific motor behavior or have different firing patterns in distinct behaviors. In mice, different groups of V0 spinal interneurons are involved in mediating left-right limb alterations at different speeds [1, 2]. In larval zebrafish, the CoLA and CoLO interneurons are specifically active in struggling and escaping behaviors, respectively [3]. In Xenopus tadpoles, a group of multifunctional commissural interneurons fires differently in swimming and struggling behaviors [4, 5]. In leech and turtles, both multifunctional and dedicated interneurons have been reported [5]. While very informative, these studies are limited by poorly characterized locomotor behaviors, lack of a comprehensive view of locomotor circuit components (MNs and PMNs) and their connectivity map, and the deficiency of genetic tools for monitoring and manipulating the activity of small population or individual neurons.

Years of work in Drosophila larvae has greatly broken through these limitations. The larval body is composed of three thoracic segments (T1-T3) and nine abdominal segments (A1-A9). Most segments contain 30 bilaterally symmetrical muscle pairs [6-8], innervated by axonal projections of around 30 glutamatergic excitatory motor neurons (MN). In the ventral nerve cord (VNC) of the CNS reside the entirety of MNs as well as interconnecting PMNs that provide input to MNs [9-12]. Drosophila larvae are capable of performing diverse motor behaviors, such forward and backward locomotion, escape rolling, turning, hunching, and head casting [13-17]. We recently used ratiometric muscle calcium imaging to characterize the muscle activity patterns underlying forward and backward locomotion in intact near free-moving animals [12]. In both modes of locomotion, waves of muscle contraction initiate from one end of the body and propagate anteriorly (forward) or posteriorly (backward), with a delay of peak activity between the sequentially recruited body segments. Both behaviors involve all muscles within each segment, but the sequence of muscle recruitment is different in forward and backward locomotion [12]. Importantly, in a collaborative work with other research groups, we recently used electron microscopy reconstruction to generate a comprehensive wiring diagram of PMNs and MNs (PMN-MN connectome) in a single larval VNC segment [10, 12, 13, 18]. This dataset has provided us an excellent opportunity to investigate how larval PMN-MN circuits generate muscle activation patterns seen in forward and backward locomotion.

One signature difference between forward and backward sequence is the recruitment timing of ventral oblique (VO) muscles 15-17, where they are recruited late during forward locomotion but become the front-runners during backward locomotion. Here, we addressed the question of how motor circuits generate these two distinct VO activity patterns. We hypothesized that the MNs innervating VO receive different neurotransmission input from PMNs during the two behaviors. By looking into the premotor circuit upstream the VO-MNs, we identified an inhibitory PMN, A06c, that shows bi-functionality in the two behaviors. A06c fires both before and after (two peaks) MNs during forward locomotion but only after MNs (one peak) during backward locomotion, restricting the activity timing of VO-MNs. A forward-dedicated excitatory PMN, A27h, fires in between of A06c’s two peak activities to recruit VO-MNs. Additionally, a forward-dedicated excitatory PMN, A18b3, forms intersegmental connection with A06c in the next anterior segment while innervating early active MNs within its own segment. Through this feedforward inhibitory motif, A18b3 activates early MNs while introducing an inhibitory delay onto VO MNs. Silencing A06c, A27h or A18b3 resulted in premature VO contraction in forward locomotion, resembling the muscle pattern in backward locomotion. Together, the A27h-A18b3-A06c circuit motif define the precise window of VO muscles’ activity timing specifically in forward locomotion, supporting our hypothesis that different muscle activity patterns can be a result of behavior-specific input from PMNs to MNs.

## Results

### Ventral oblique (VO) muscles show different activity timing during forward and backward locomotion

The peristaltic motor activity within the 12 body segments in larvae moves from posterior to anterior segments during forward, and vice versa during backward locomotion (**Figure 1A-C**). Calcium imaging of individual muscles has revealed that all body wall muscle types are activated in both forward and backward locomotion; however, a subset of muscles show different activity timing between these two behaviors. Muscles 15, 16 and 17, hereafter named the ventral oblique (VO) muscles, are examples of such differentially activated muscle groups [12]. During backward locomotion, VO muscles in a given segment are the earliest activated muscles, while during forward locomotion they are activated significantly later than many other muscles. To corroborate this interesting finding, we used confocal microscopy combined with muscle calcium imaging to examine the activation patterns of VO muscles, relative to others, during forward and backward locomotion in intact animals. Our data further confirmed that VO muscles indeed contract earlier in backward than in forward locomotion (**Figure 1D; Video 1**). This data provides an excellent system to understand how the same set of MNs/Muscles are activated in different patterns to generate distinct motor outputs (i.e., forward and backward locomotion). With an emphasis on VO muscles, we address this fundamental question in the following sections.

**Figure 1.**
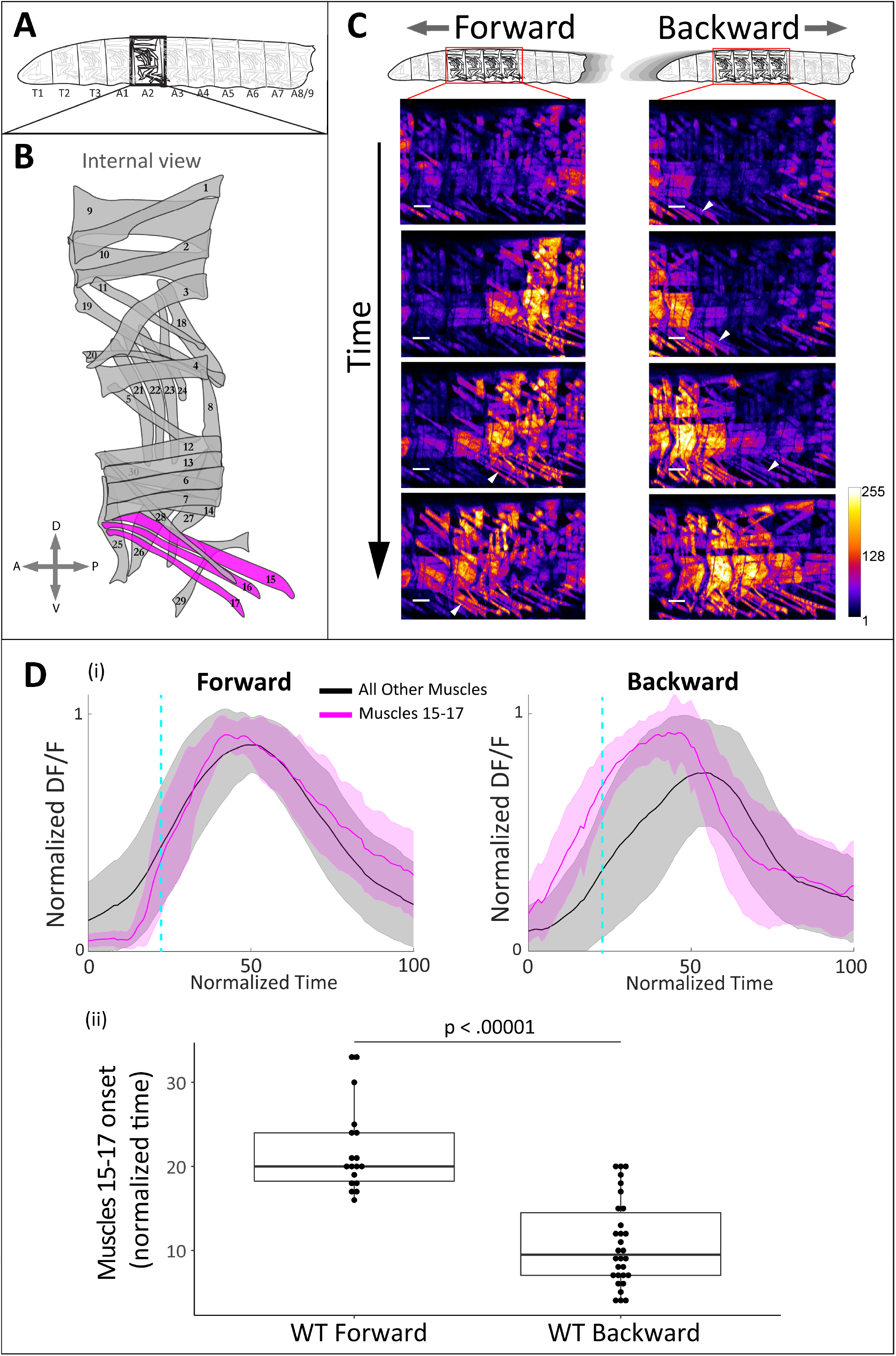
The ventral oblique (VO) muscles are recruited earlier in backward than in forward locomotion. **A)** The larval body consists of 3 thoracic and 9 abdominal segments. Segments A1-A7 share a similar musculature as shown in B). **B)** Each segment is bilaterally symmetric with around 30 body wall muscles in each hemisegment. Muscles 15, 16 and 17 (“VO muscles”; magenta) are the focus of this study. **C)** Sequential still images of muscle GCaMP6f signal during wild type forward (left) and backward (right) locomotion. Scale bar: 50μm. Arrowheads show VO muscles of a given segment being recruited. Genotype: GMR44H10-LexA lexAOP-GCaMP6f; lexAOP–mCherry. **D)** Within a segment, VO muscles are recruited earlier in backward than forward crawling. i) Diagrams showing GCaMP6f of VO muscles (magenta) and all other muscles (black) within each segment during a forward or backward crawl bout, normalized to mCherry. Normalized time 0-100 represents the start and end of a crawl cycle standardized based on the comprehensive activity of the entire segment using a previously described PCA-based method (Zarin et al. 2019). Dashed line represents t = 25 normalized time. Error bar (shaded area) represents standard deviation. n = 25 crawl bouts in 5 animals for forward and n = 19 bouts in 4 animals for backward. ii) The activity onset points of VO muscles were earlier in backward than in forward crawling. Onset time is defined as the time point where the muscle GCaMP6f activity first reaches 20% of its peak activity during a crawl. Muscle data reused from i). Student’s t test, n = 18 individual muscles in forward and 30 in backward p<0.00001.

### VO MNs receive highly specific excitatory and inhibitory premotor inputs

Larval muscles executing locomotion are innervated by two types of glutamatergic excitatory MNs: Is MNs (forming <u>s</u>mall boutons) and Ib MNs (forming <u>b</u>ig boutons) [19-26]. VO muscles 15-17 are innervated by two pairs of Ib (MN15/16 and MN15/16/17, hereafter named the VO MNs) and a pair of Is MNs (**Figure 2A**). While VO MNs specifically innervate VO muscles, type Is MN innervates multiple muscles including two of the VO muscles (**Figure 2A**). Given their VO specific innervation, we reasoned that VO MNs (MN15/16 and MN15/16/17) should underlie distinct activity patterns of VO muscles in forward and backward locomotion. We thus sought to identify the individual premotor neurons (PMNs) that may regulate these MNs. We used our recently published premotor-motor neuron (PMN-MN) connectome to look for PMNs that heavily synapses onto VO MNs, and identified two PMN pairs (A06c and A27h) as possible candidates (**Figure 2A, B**). A06c is a GABAergic inhibitory PMN that almost exclusively synapses onto MN15/16 and MN15/16/17, while A27h is a cholinergic excitatory PMN that makes a high number of synapses with VO MNs and fewer synapses with Is MN (**Figure 2B**). Interestingly, both A06c and A27h PMNs are bilaterally connected to their downstream MNs, meaning that A06c and A27h in a given hemisegment directly connect to all MNs located in both left and right hemisegments (Figure 2B, C). Strikingly, A27h also establishes direct synapses onto A06c within its own segment, resulting in an intra-segmental feedforward inhibitory PMN-PMN-MN motif (**Figure 2A**). Other than A06c and A27h, several other PMNs (such as A18b2, A02i, A02g) directly synapse onto VO-MNs (**Figure 2B**). These PMNs, however, establish unilateral and significantly lower number of synapses with VO-MNs as well as several other MNs (Figure 2B, C). Unilateral refers to a connectivity pattern where a PMN in a given hemisegment connects to either left or right but not both counterparts of each MN pair (**Figure 2C**). Taken together, analysis of PMN-MN connectome suggested that A27h and A06c PMNs may be crucial for differential activity of VO-MNs in different behaviors. We further investigate this possibility in the following sections.

**Figure 2.**
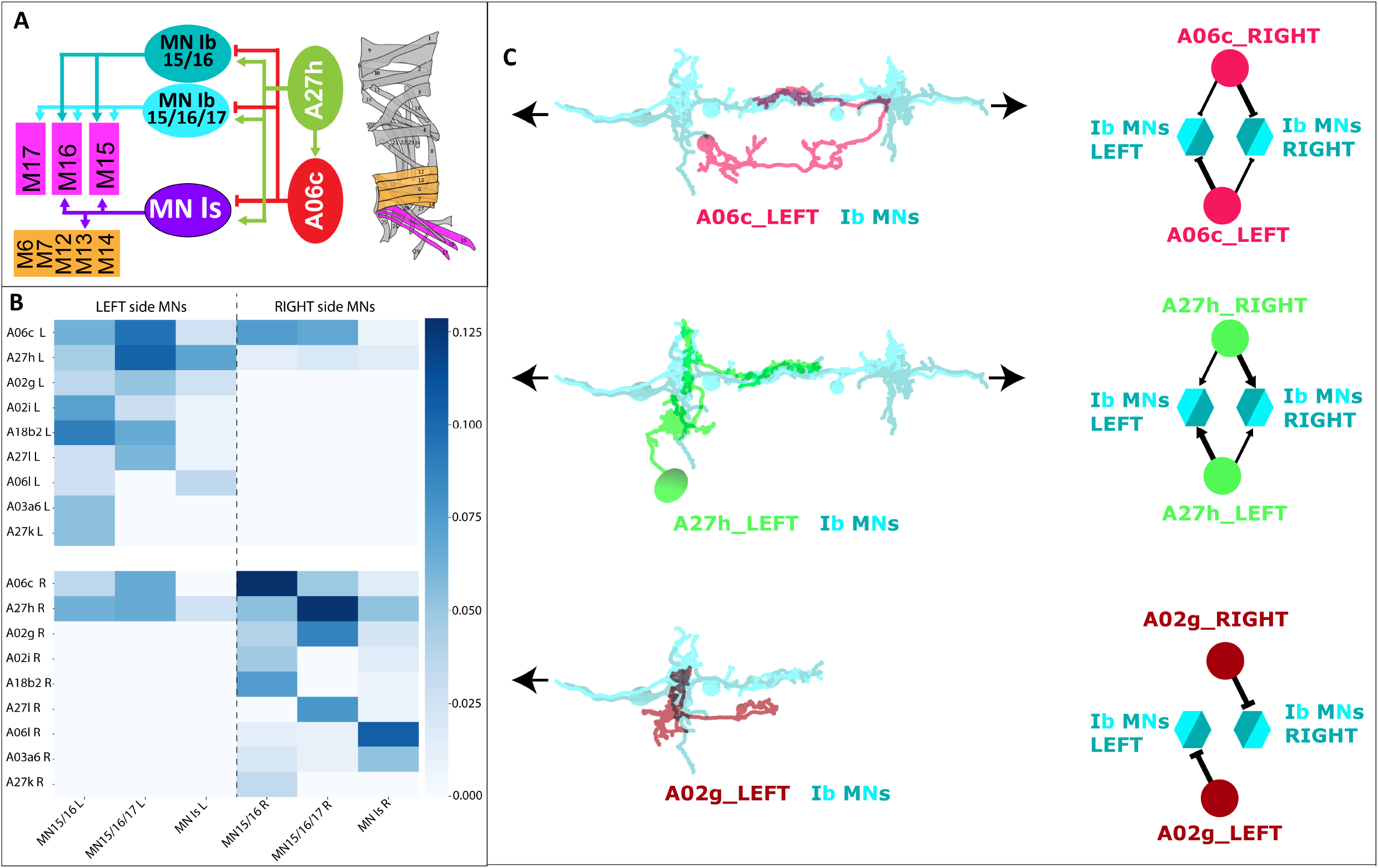
The VO MNs receive inputs from excitatory and inhibitory PMNs. **A)** Two type Ib MNs exclusively innervate VO muscles (magenta), while a type Is MN innervate VO muscles as well other ventral muscles (orange). Both type Ib and Is MN receive inputs from A27h (excitatory-green) and A06c (inhibitory-red) PMNs. **B)** A heatmap showing the connectivity patterns between PMNs (rows) and three different VO MNs. MNs and PMNs have been divided based on left (L) and right (R) hemisegments. A27h and A06c are two main PMNs providing inputs to these MNs. **C)** A06c and A27h show bilateral connectivity patterns with MNs (i.e., a given PMN in left or right hemisegment connects to MNs in both left and right hemisegment). A02g, in contrast, is an example of a unilateral PMN where a given PMN in left or right hemisegment connects to MNs in only left or right hemisegment.

### A06c and A27h provide behavior-specific inputs onto VO MNs

We next determined if A27h and A06c PMNs have distinct activity patterns in forward and backward locomotion. We expressed GCaMP6m in a subset of type Ib MNs (MN2, 3, 4, 9, 10; hereafter named the reference MNs) and jRCaMP1b in A06c or A27h PMNs and recorded fictive forward and backward locomotion in isolated brains. A perfectly isolated larval brain frequently exhibits spontaneous forward and backward fictive crawls, wherein motor activity propagates from posterior to anterior VNC segments in forward and vice versa in backward locomotion.

We first investigated the activity pattern of A06c PMN. During forward locomotion, A06c in a given segment has two subsequent peaks of activity, respectively before and after the peak activity of the reference MNs within that segment (**Figure 3A; Video 2**). Interestingly, during backward locomotion, A06c is only active after the reference MNs (**Figure 3A; Video 2**). The lack of first inhibitory peak of A06c PMN in backward locomotion suggests that VO-Ib-MNs do not receive any early inhibitory inputs in backward, which is in turn consistent with the earlier activation of VO muscles in backward than in forward locomotion. Based on A06c’s different activity patterns in two behaviors, we name A06c PMN a “bi-modal PMN”.

**Figure 3.**
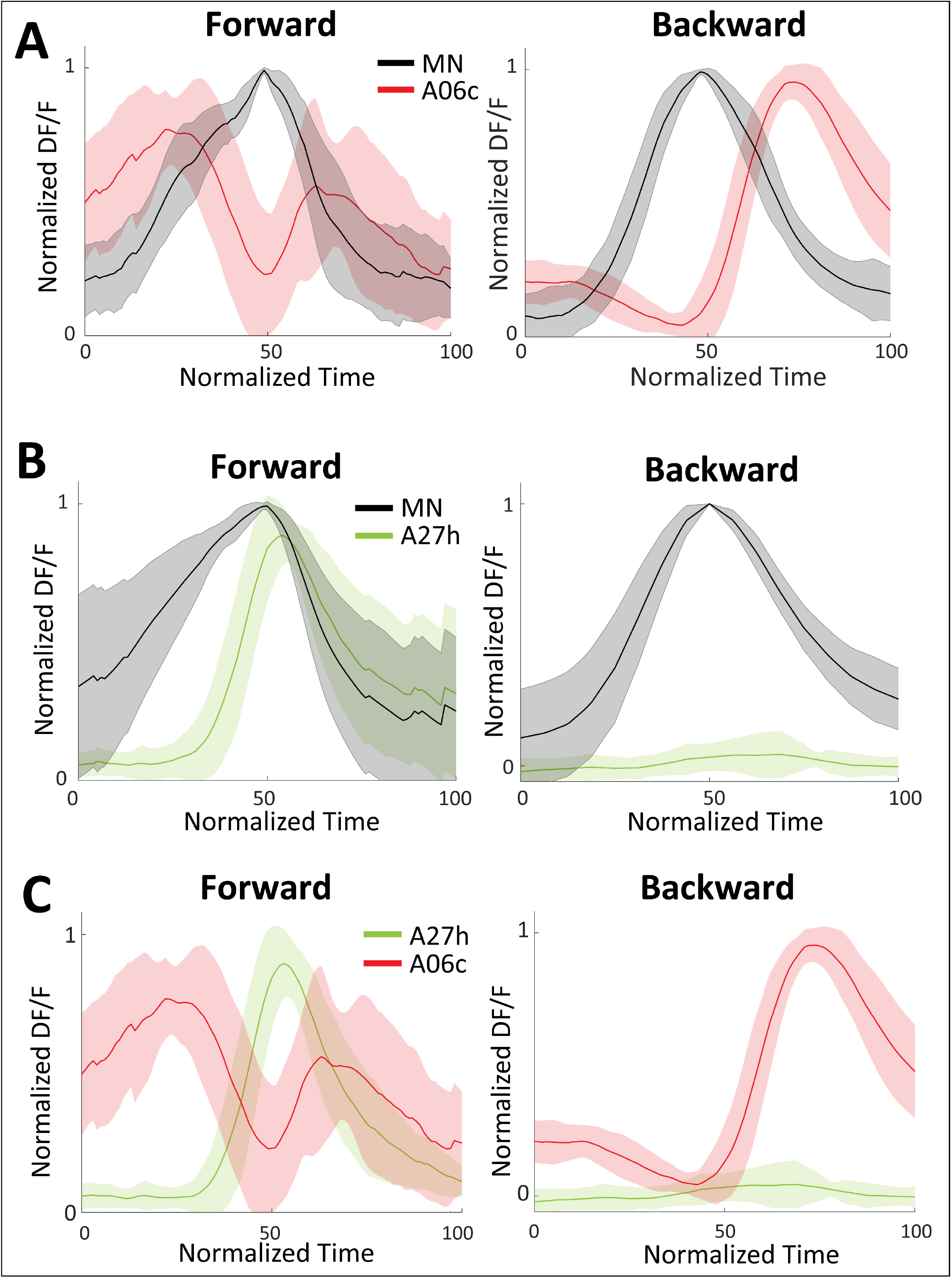
A06c is a bi-modal and A27h is a forward-dedicated PMN. **A)** Activity of A06c relative to reference MNs within the same segment during fictive forward and backward locomotion. A06c showed two peaks of activity, both before and after MNs, during forward locomotion, but only one peak (after MNs) during backward locomotion. Data from different brains was normalized and aligned based on MN activity. n = 18 fictive crawls from 5 brains. Genotype: A06c-split-Gal4 > UAS-jRCaMP1b; CQ-LexA > LexAOP-GCaMP6m. **B)** Activity of A27h (magenta) relative to MN (black) during forward and backward locomotion. A27h only fires during forward locomotion, after the reference MNs. n = 7 fictive crawls from 3 brains. Genotype: R36G02-Gal4 (A27h) > UAS-jRCaMP1b; CQ-LexA > LexAOP-GCaMP6m. **C)** Merged graphs showing activity of A06c, A27h and MN during forward and backward locomotion. In forward locomotion, A27h is active in-between A06c’s first and second peak. Data from different genotypes was aligned and merged based on GCaMP6m (MN) activity. In all panels, error bars (shaded area) represent standard deviation. Part of the data used in panel C has been previously published in another paper published by the same author [12].

We then examined the activity pattern of the excitatory A27h PMN in forward and backward fictive locomotion. During forward locomotion, A27h excitatory PMN begins to fire slightly after the reference MNs, reaching its peak activity in a time window between A06c’s first and second peak (**Figure 3B-C; Video 3**). This data combined with A27h-A06c-MN connectome suggests that, during forward locomotion, A06c defines a temporal window during which A27h provides excitatory inputs onto VO-MNs. As reported before [12, 18], A27h has no activity during backward locomotion, suggesting that A27h excitatory input is not involved in activation of VO-MNs during backward locomotion. Because A27h is exclusive to forward locomotion, we name it a “forward-dedicated PMN”.

### A06c and A27h Loss of function leads to premature VO recruitment in forward locomotion

We reasoned that the bi-modal inhibitory A06c and forward-dedicated excitatory A27h PMNs provide behavior-specific inputs onto VO-MNs, thereby leading to distinct VO muscle activity in forward and backward locomotion. To test this, we silenced A06c or A27h neurons and determined the effect on larval forward locomotion in intact behaving animals. We silenced the neurons of interest with two different approaches. For permanent neuronal silencing, we used inward rectifying Potassium channel (Kir2.1); for temporary silencing, we used the light-gated *Guillardia theta* anion-conducting channelrhodopsin-1 (GtACR1) [27](**Figure 4A**).

**Figure 4.**
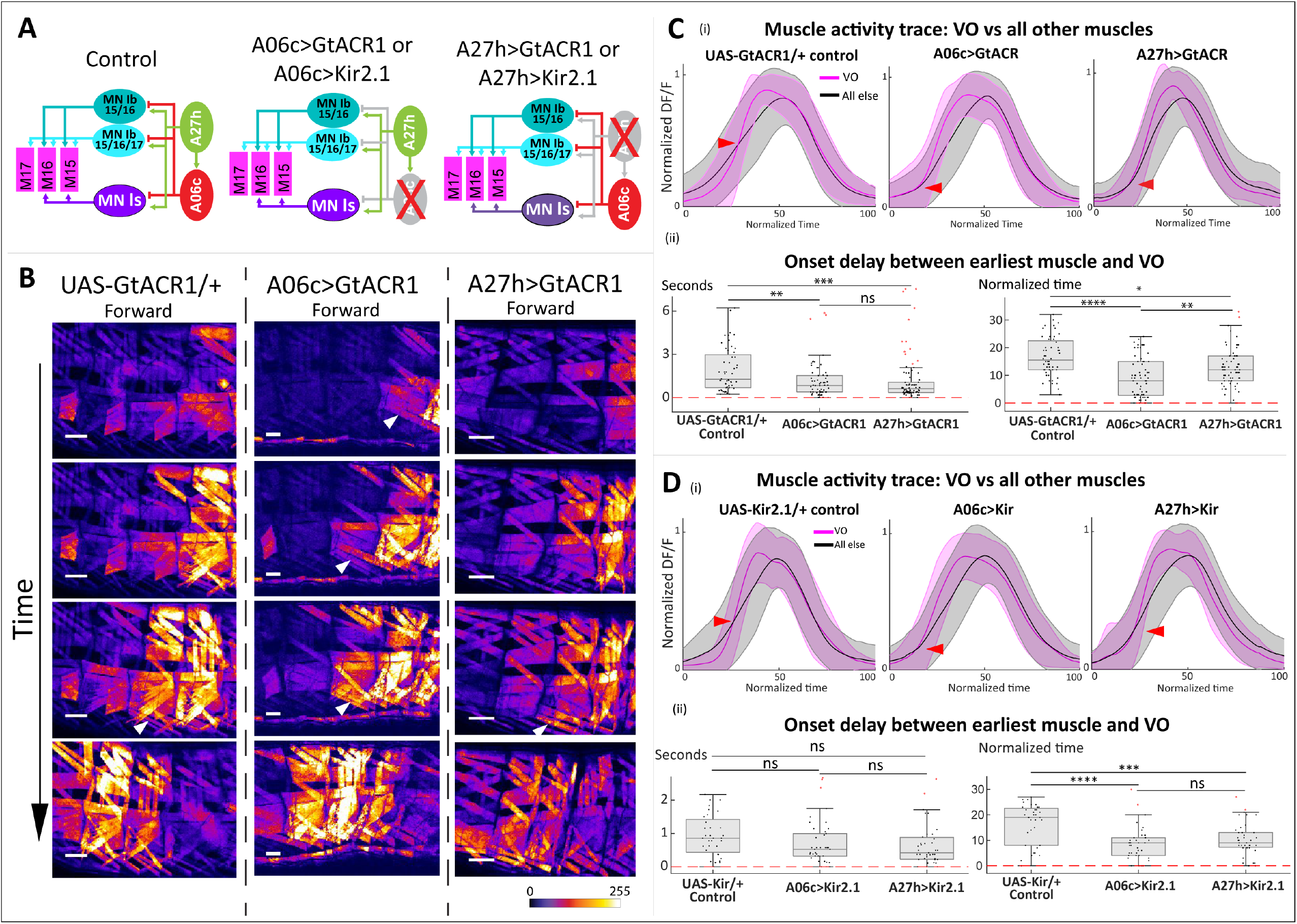
A06c and A27h loss of function (LOF) results in premature VO activity during forward locomotion. **A)** Schematics of performed experiments. **B)** Still images showing a forward locomotion trial in control animals, and A06c or A27h silenced animals. Arrowheads point out contracting VO muscles. In A06c silenced animals, VO muscles in each segment were prematurely recruited. Arrowheads indicate contracting VO muscles. Scale bars: 50 μm. **C, D)** Quantification of muscle GCaMP activity and onset timings showed that VO muscles were recruited early in A06c LOF and A27h LOF animals. **C, D (i))** Line plots showing GCaMP activity of VOs compared to all other muscles within the segment. In animals where A06c or A27h PMNs were silenced using either GtACR1 or Kir2.1, VO muscles were recruited earlier than other muscles within the same segment. For each genotype, individual muscles were quantified, and different trials were aligned and normalized based on the peak activity of the entire segment. n = 27 crawl bouts in 10 animals for UAS-GtACR1 control, 22 crawl bouts in 7 animals for A06c>GtACR1, 18 crawl bouts in 5 animals for A27h>GtACR1, 17 crawl bouts in 5 animals for UAS-Kir2.1 control, 22 bouts in 8 animals for A06c>Kir2.1, 18 bouts in 6 animals for A27h>Kir2.1. **C, D (ii))** During forward locomotion, silencing A06c or A27h PMN leads to significant decrease in the temporal delay between VO and the earliest muscle within the same segment. Boxplots: delay of onset timings between individual VO muscles (15, 16 or 17) and the earliest recruited muscle within the segment, in seconds (left) or normalized time (right). Onset time defined as the time point (in seconds or in normalized time) when a muscle’s activity reaches 20% of the maximum increment before peaking. Data points represent individual muscles’ timings. One-way ANOVA with Tukey HSD post-hoc test (for GtACR1 loss of function: normalized time, F = 15.1928, p < .00001, time in seconds, F = 7.6533, p = .000638; for Kir2.1 loss of function: normalized time, F = 13.53003, p < .00001, time in seconds, F = 1.81559, p = .167415). p values for pair-wise comparisons: A06c>GtACR1 vs UAS-GtACR1/+ control: normalized time, p < .00001, time in seconds, p = .00251, A27h>GtACR1 vs UAS-GtACR1/+ control: normalized time, p = 0.045, time in seconds, p = .00090; A06c>GtACR1 vs A27h>GtACR1: normalized time: p = .00370, time in seconds, p = .95346; A06c>Kir2.1 vs UAS-Kir2.1/+ control: normalized time, p = .00001, time in seconds, p = .20150, A27h>Kir2.1 vs UAS-Kir2.1/+ control: normalized time p = .00050, time in seconds, p = .23460; A06c>Kir2.1 vs A27h>Kir2.1: normalized time: p = .49360, time in seconds, p = .99600.. Genotypes: A06c-split-Gal4 > UAS-GtACR1.d.EYFP; 44H10::GCaMP6f (“A06c>GtACR1”), A06c-split-Gal4 > UAS-Kir2.1-GFP; 44H10::GCaMP6f (“A06c>Kir2.1”), 36G02-Gal4 (A27h) > UAS-GtACR1.d.EYFP; 44H10::GCaMP6f (“A27h>GtACR1”), R36G02-Gal4 (A27h) > UAS-Kir2.1-GFP; 44H10::GCaMP6f (“A27h>Kir2.1”), UAS-Kir2.1-GFP/+; 44H10::GCaMP6f (“UAS-Kir2.1/+ Control”) and UAS-GtACR1.d.EYFP/+; 44H10::GCaMP6f (“UAS-GtACR1/+ Control”). 36G02-Gal4 line targets A27h as well as another neuron known as A03g or M neuron [46, 55]. However, A03g/M neuron does not make any synapses with VO MNs.

We hypothesized that A06c’s first activity peak introduces an inhibitory delay onto VO MNs during forward locomotion, and thus A06c loss of function should cause premature VO muscle activity. Indeed, quantification of muscle GCaMP imaging showed that, in animals where A06c was silenced with either GtACR1 or Kir2.1, VO muscles were recruited prematurely during forward locomotion (**Figure 4 B-D; Video 4**). These results, together with our dual-color calcium data, suggest that A06c PMN provides early inhibition onto VO MNs to block premature firing, maintain the precise recruitment timing, generate forward-specific VO muscle pattern, and therefore ensure the orderly and effective recruitment of muscles.

We next examined A27h’s role in VO muscle activation. We hypothesized that after A06c’s first peak activity during forward locomotion, A27h provide excitation to recruit VO muscles. As one of the major sources of excitatory input onto its target MNs, we expected that silencing of A27h might lead to lowered excitability of these MNs and thus abnormal timing of muscle recruitment, or even failure in recruiting VO muscles during forward locomotion. Surprisingly, in A27h loss-of-function animals, VO muscles were still recruited during forward locomotion albeit with a premature onset compared to control animals. (**Figure 4B-D; Video 5**). This could be explained by the possibility that in the absence of A27h inputs, VO MNs receive enough excitation from other excitatory PMNs and are able to activate VO muscles. In fact, based on connectome data, while A27h is the major excitation source for these Ib MNs, it is only one of the several excitatory PMNs that provide excitation type Is MN. However, silencing A27h impairs precise integration of Ib and Is inputs onto VOs, thereby resulting in abnormal (premature) VO activity. Taken together, our results showed that A06c indeed provides early inhibition to delay VO activity during forward locomotion; excitation from A27h to VO MNs is not necessary for VO recruitment but is necessary for precise timing of VO recruitment.

### The excitatory A18b3 PMN drives A06c-mediated inhibition of VO muscles in forward locomotion

A06c fires both before and after motor neurons in forward locomotion, while it fires only after motor neurons in backward locomotion. What is the circuit mechanism underlying A06c’s forward-specific first peak activity? To address this, we went back to our TEM dataset and, using A06c in segment A1 as ground-truth, reconstructed A06c PMNs as well as their pre-synaptic partners in two additional segments (T3 and A2). We analyzed the connectome and identified a cholinergic excitatory PMN, A18b3, that makes a significant number of synapses onto A06c of the next anterior segment (**Figure 5 A-B**). A18b3 also synapses with two early firing type Ib MNs (MN Ib 14 and MN Ib 30) in its own segment (data not shown). Based on its morphology, we found A18b3 to be identical to a previously studied neuron known as <u>c</u>holinergic lateral interneuron 1 (CLI1) [28] (**Figure 5, figure supplement 1)**. Based on that study, CLI1 is a forward-dedicated PMN; i.e., active only during forward but not backward locomotion [28]. Therefore, while activating early MNs/Muscles (M14 and M30) in its own segment, A18b3 (CLI1) could also inhibit and delay VO muscles of the next anterior segment via generating A06c’s forward-specific first peak. To test this, we performed dual-color calcium imaging to determine the firing pattern of A18b3 and its phase relationship with A06c. Our result confirmed that, as reported before, A18b3 is dedicated to forward locomotion; most importantly, A18b3 fires in sync with the first peak of A06c PMN located in the next anterior segment (**Figure 5C; Video 6**). These data support our hypothesis that A18b3 provides feedforward inhibition on VO MNs via A06c, thereby delaying activation of VO muscles during forward locomotion.

**Figure 5.**
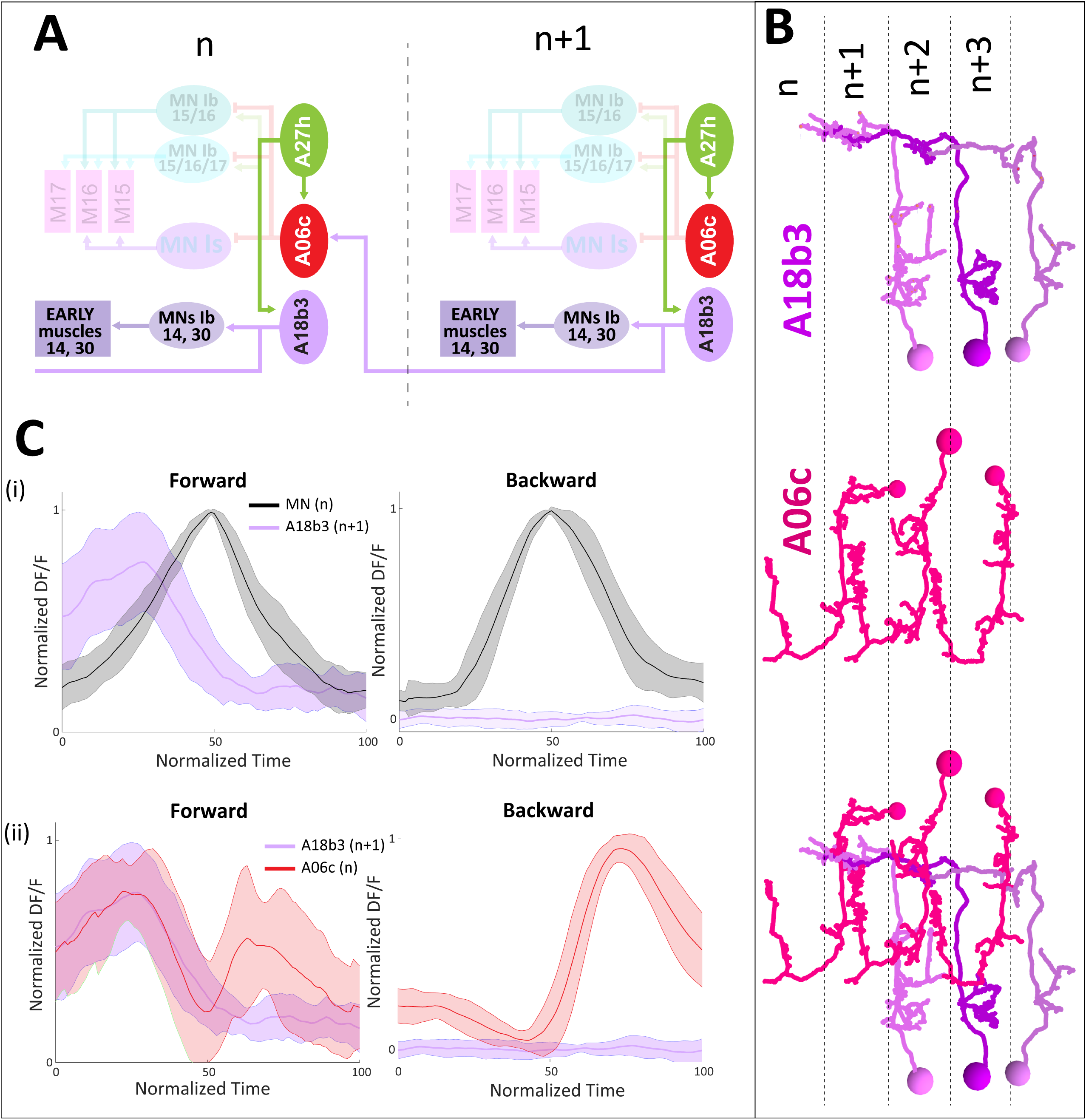
A forward-dedicated excitatory PMN, A18b3, fires in synchronous with the first activity of A06c of the next anterior segment. **A)** diagram showing intersegmental connection between A18b3 (segment n+1) and A06c of the next anterior segment (segment n). A18b3 also innervate early forward MNs, while receiving input from A27h of the same segment. **B)** TEM reconstruction of A18b3 (purple) and A06c (red) in four subsequent VNC segments. A18b3 projects to the next anterior segment where it synapses with A06c. **C)** A18b3 is a forward dedicated PMN, and fires in synchronous with the first peak of activity of its target A06c. i) A18b3 (n+1) fires before the reference MNs of the next anterior segment (n) and is silent during backward locomotion. n = 19 fictive crawls in 5 animals for forward, 18 fictive crawls in 5 animals for backward. ii) A18b3 activity during forward locomotion overlaps with the first peak of its target A06c. A27h data reused from figure 3. Shaded areas represent standard deviation. Genotype: A18b3-split-Gal4 > UAS-jRCaMP1b; CQ-LexA > LexAOP-GCaMP6m.

Next, we reasoned that if A18b3 indeed causes A06c’s first peak, silencing A18b3 should phenocopy the defect caused by A06c loss of function; i.e., premature recruitment of VO muscles during forward locomotion. To test this, we silenced A18b3 (with Kir2.1 or GtACR1) and studied the effect on animals performing forward locomotion using muscle GCaMP imaging (**Figure 6A-B**). Indeed, similar to A06c loss of function, silencing A18b3 led to significant premature VO contraction, where the delay between VO and the earliest muscles within the segment was greatly decreased or almost diminished, (**Figure 6 C-D; Video 7**). The premature activation of VO muscles during forward locomotion of animals lacking A06c or A18b3 is reminiscent of early VO activation in backward locomotion, wherein there is no A06c’ initial peak or A18b3 activity. A18b3 represents an excitatory PMN that exerts antagonistic input onto different MNs, thereby generating an activation order for muscles.

**Figure 6.**
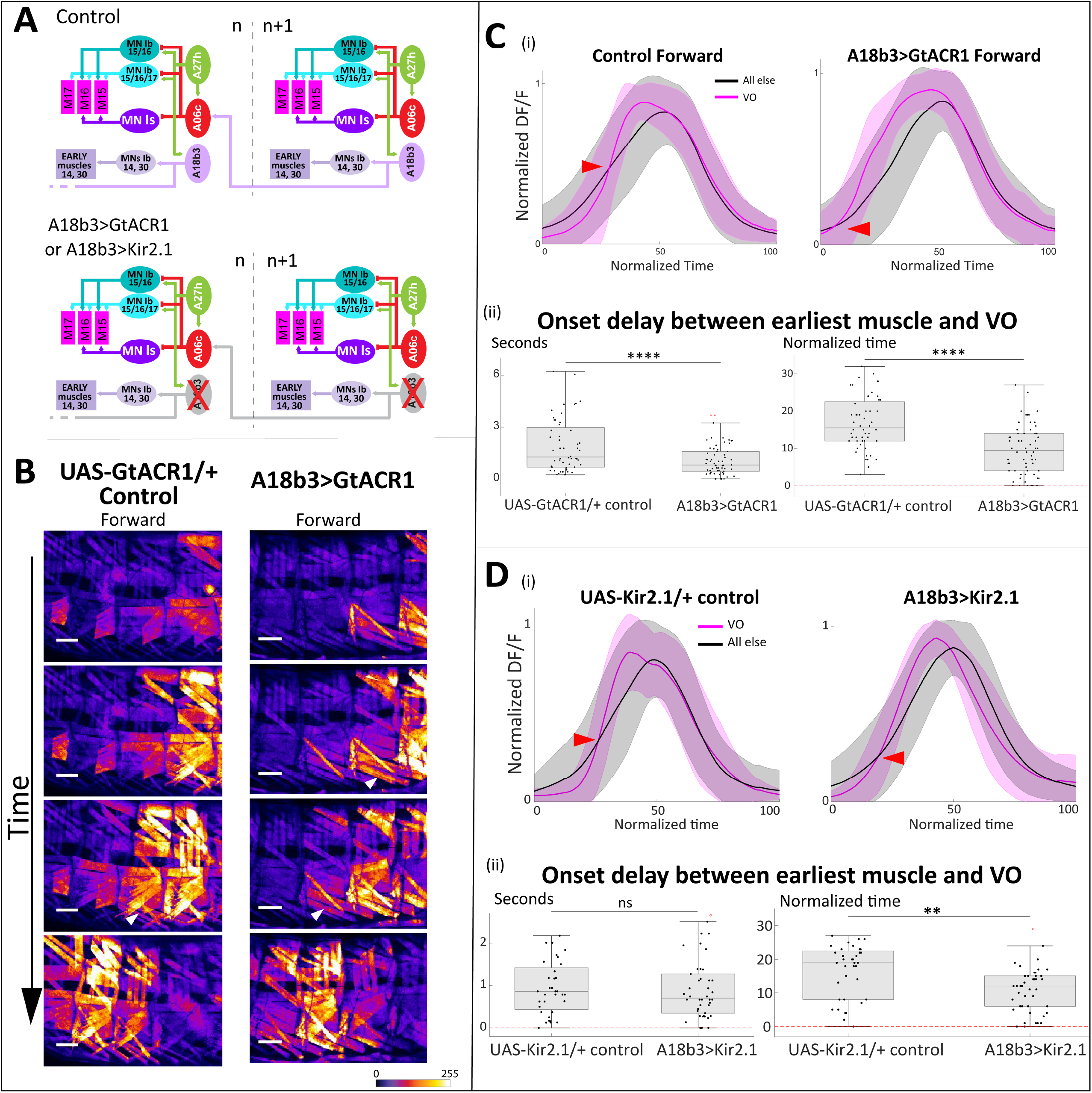
A18b3 loss of function results in premature VO contraction during forward locomotion. **A)** schematics of performed experiments. **B)** Still images showing a forward locomotion trial in UAS-GtACR1/+ control and A18b3>GtACR1 animals. Arrowheads point out contracting VO muscles. **C, D (i))** Line plots showing GCaMP activity of VOs compared to all other muscles within the segment. A18b3 LOF animals showed premature VO contraction. n = 22 crawl bouts in 9 animals for A18b3>GtACR1 and 18 crawl bouts in 6 animals for A18b3>Kir2.1 animals. **C**,**D (ii))** During forward crawl, the delay between VO and the earliest muscle was substantially decreased in A18b3 LOF animals compared to control. Boxplots: delay of onset timings between individual VO muscles (15, 16 or 17) and the earliest recruited muscle within the segment, in seconds or normalized time. Data points represent individual muscles’ timings (reused from C) i) and D) i)). T-tests p values: for GtACR1 loss of function: normalized time, p < .00001, time in seconds, p < .00001; for Kir2.1 loss of function: normalized time, p = .000541, time in seconds, p = .325216. Genotypes: A18b3-split-Gal4 > UAS-GtACR1.d.EYFP; 44H10::GCaMP6f (“A18b3>GtACR1”) and A18b3-split-Gal4 > UAS-Kir2.1-GFP; 44H10::GCaMP6f (“A18b3>Kir2.1”).

Here we summarize the sequence of events occurring between different PMN-MN circuit components to ensure that VO muscles are activated at the right time during forward locomotion (**Figure 7**). Our findings provide important insight onto how the differential function of a subset of dedicated or bi-modal PMNs sculpts MN dynamics, enabling them to be multifunctional and generate diverse motor outputs. Evolution of dedicated and bi-modal PMNs seem to be cost-effective, exempting the animals from the need to have a dedicated set of MNs/muscles for different behaviors (e.g., larval forward and backward locomotion).

**Figure 7.**
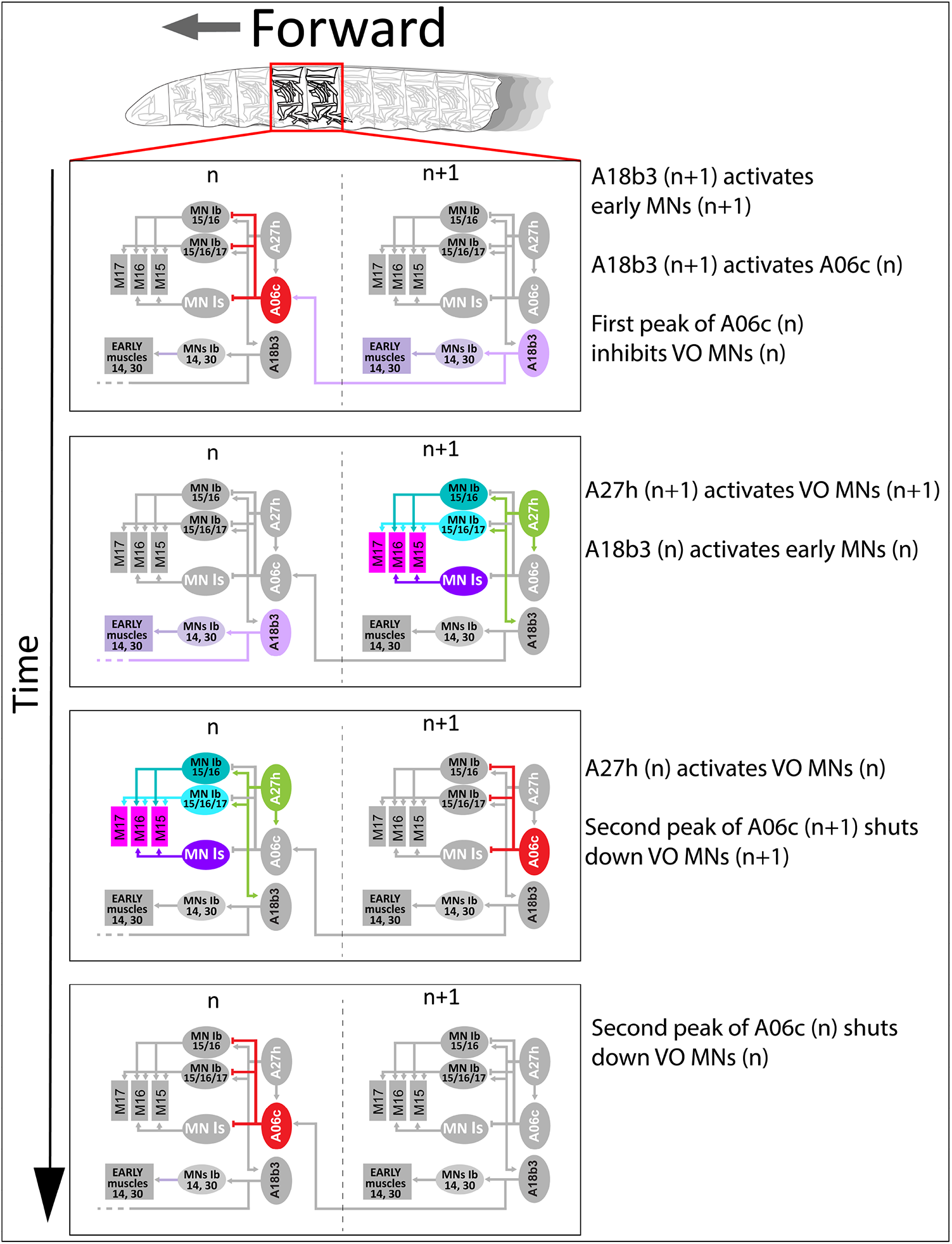
A model describing how A27h-A18b3-A06c motif defines activation window of VO muscles during forward locomotion.

## Discussion

The ability to generate different locomotor behaviors is essential to survive in an ever-changing living habitat. Studies of different animal models have provided valuable insights on how motor circuits produce diverse motor outputs and how different muscles are coordinated during execution of a given motor behavior. In leech, some PMNs rhythmically oscillate in both swimming and crawling but with different frequencies; some fire in crawling but are silent in swimming [29]. In marine mollusk Tritonia Diomedea, the dorsal swim interneurons also make direct connections with the crawling-related cilia MNs and can induce crawling [30]. In vertebrates such as turtle, zebrafish and mouse, groups of spinal premotor INs have been characterized based on their specific functions in locomotion [5, 31, 32]. In zebrafish, different groups of V2a interneurons are associated with slow, medium or fast MNs have different activity pattern during swimming, ensuring the orderly recruitment of the MNs/muscles [33, 34]. In tetrapod, coordination of different limbs may be mediated by specific cluster of spinal interneurons [35, 36], such as the commissural V0d interneurons in mice that are responsible for left-right coordination in walking gait [32, 37, 38]. Multiple studies have demonstrated the “size principle” of the coordination of different muscles/MNs in a single behavior in numerous organisms from insects to human, indicating that small MNs (controlling small muscles and slow movements) are recruited prior to large MNs (controlling large muscles and fast movements)[39-44]. While very informative, many of these studies were not able to reach deep into the neural mechanism underlying multifunctionality of motor circuits, due to poor spatial and temporal resolution, lack of comprehensive view of circuit connectivity and lack of genetic approaches to monitor individual cells. Our study took advantage of the recently built larval locomotor circuit connectome [10, 12, 18, 45-47] and the rich collection of genetic and optogenetic tools [48, 49] to revisit and address this question in intact freely moving Drosophila larvae at a single-muscle and single-neuron resolution.

Here, we reveal a circuit mechanism for how behavior-specific function of PMNs leads to behavior-specific muscle activation. We show that an inhibitory PMN, A06c, connects to two excitatory PMNs, A27h and A18b3 to form a feedforward intersegmental circuit motif that exclusively functions in forward locomotion to precisely regulate the contraction timing of VO muscles (**Figure 7**). This motif introduces an inhibitory delaying input onto VO MNs only during forward locomotion, thereby leading to a delayed contraction of VO muscles. Such inhibitory input onto VO muscles is absent in backward locomotion, resulting in earlier VO muscle activation in backward than in forward locomotion. Our result provide insight into the neural mechanism of motor pattern generation in different behaviors. It highlights that PMNs, in addition to monosynaptic connections with MNs, establish intricate connections with each other to produce functionally significant circuit motifs. An organism as simple as Drosophila larva, uses these PMN-PMN motifs to (i) exert antagonistic effects on downstream MNs, (ii) prevent premature activation of MNs, (iii) restrict duration of excitatory inputs onto MNs, and (iv) shut down the MNs once their activity is no longer needed. Our work supports the notion that EM reconstruction of neuroanatomical connectome is necessary but not sufficient to fully understand how neural circuits works [50]. The discovery of bi-modal and dedicated PMNs clearly demonstrate that a given neuroanatomical circuit may not necessarily act the same way during different behaviors. Thus, to achieve a comprehensive understanding of any circuit works to contribute to a given behavior, both functional and neuroanatomical examination of its components is necessary.

## Materials and methods

### Key resources table

| Reagent type (species) or resource | Designation | Source or reference | Identifiers | Additional Information |
| --- | --- | --- | --- | --- |
| Genetic reagent (*Drosophila melanogaster*) | R36G02-Gal4 | Bloomington Drosophila Stock Center | BDSC:49939 | A27h line |
| Genetic reagent (*Drosophila melanogaster*) | A06c split Gal4 | This paper | SS21783 | A06c split line |
| Genetic reagent (*Drosophila melanogaster*) | A18b3 split Gal4 | This paper | SS04114 | A18b3 split line |
| Genetic reagent (*Drosophila melanogaster*) | 44H10::GCaMP6f | This paper |  | 44H10 is a muscle enhancer element. This fly stock expresses GCaMP6f in all larval muscles |
| Genetic reagent (*Drosophila melanogaster*) | 44H10-LexA | Bloomington Drosophila Stock Center | BDSC:61543 | This fly stock expresses LexA in larval muscles |
| Genetic reagent (*Drosophila melanogaster*) | CQ-LexA | Gift from Dr. Chris Doe |  | CQ is an enhancer element that targets reference motor neurons: MN2, MN3, MN4, MN9, MN10. This line expresses Lex-A in CQ MNs. |
| Genetic reagent (*Drosophila melanogaster*) | LexAop-GCaMP6m | Bloomington Drosophila Stock Center | BDSC:44275 | This fly stock expresses GCaMP6m under the control of LexA operator. GCaMP6m is a green calcium indicator that is compatible with jRCaMP1b |
| Genetic reagent (Drosophila melanogaster) | LexAop-GCaMP6f | Bloomington Drosophila Stock Center | BDSC:44277 | This fly stock expresses GCaMP6m under the control of LexA operator. |
| Genetic reagent (Drosophila melanogaster) | UAS-jRCaMP1b | Bloomington Drosophila Stock Center | BDSC:63793 | This fly stock expresses jRCaMP1b under the control of UAS. jRCaMP1b is a red Calcium indicator compatible with GCaMP6m |
| Genetic reagent (Drosophila melanogaster) | UAS-GtACR1.d.EYFP | Bloomington Drosophila Stock Center | BDSC:92983 | Expresses light-gated anion channel GtACR1 tagged with EYFP under the control of UAS. GtACR1 is an optogenetic tool that, once exposed to blue and green light, causes membrane hyperpolarization and inhibits neuron activity. |
| Genetic reagent (Drosophila melanogaster) | UAS-Kir2.1-EGFP | Bloomington Drosophila Stock Center | BDSC:6595 | Expresses human KCNJ2 (Kir2.1) tagged with EGFP under the control of UAS. Kir2.1 is an inwardly-rectifying potassium channel that leads to hyperpolarization of neurons. |
| Genetic reagent (Drosophila melanogaster) | 44H10::GCaMP6f, UAS-GtACR1.d.EYFP | This paper |  | 44H10::GCaMP6f recombined with UAS-GtACR1.d.EYFP on 3<sup>rd</sup> chromosome |
| Genetic reagent (Drosophila melanogaster) | 44H10::GCaMP6f, UAS-Kir2.1-EGFP | This paper |  | 44H10::GCaMP6f recombined with UAS-Kir2.1-EGFP on 3<sup>rd</sup> chromosome |
| Genetic reagent (Drosophila melanogaster) | 44H10-LexA, LexAop-GCaMP6f | [12] |  | Fly stock expressing GCaMP6f in larval muscles. 44H10-LexA and LexAop-GCaMP6f recombined on 2<sup>nd</sup> chromosome. |
| Genetic reagent (Drosophila melanogaster) | GMR44H10-LexA LexAOP-GCaMP6f; LexAOP-mCherry | [12] |  | Fly stock expressing GCaMP6f and mCherry in larval muscles. |
| Genetic reagent (Drosophila melanogaster) | LexAop-GCaMP6m, UAS-jRCaMP1b | This paper |  | LexAop-GCaMP6m and UAS-jRCaMP1b recombined on 3<sup>rd</sup> chromosome. |
| Genetic reagent (Drosophila melanogaster) | CQ-LexA; LexAop-GCaMP6m, UAS-jRCaMP1b | This paper |  | Fly stock carrying three different elements. CQ-LexA is on 2<sup>nd</sup> chromosome. LexAop-GCaMP6m and UAS-jRCaMP1b recombined on 3<sup>rd</sup> chromosome. |
| chemical compound, drug, chromophore | all trans-Retinal | Millipore Sigma-Aldrich | R2500-100MG | A chromophore molecule that binds to GtACR1 and other opsin channels, act as a co-factor, and sensitizes them to light exposure in optogenetic experiments. |
| Software, algorithm | MATLAB R2021b | The MathWorks, Inc. |  |  |
| Programming language, software, algorithm | Python Programming Language | Python Software Foundation | Used for analyzing the TEM connectome | free and open-source software for computing and graphics |
| Software | Rstudio | Rstudio |  | free and open-source software for statistical computing and graphics |
| Software, Data Science Platform | Anaconda |  | Used for executing Python scripts | An open-source distribution for python and R languages |
| Software | G*Power 3.1 | [51] | G*Power 3.1 | A free to use software to compute statistical power analyses for different tests |
| Microscope software | ZEISS ZEN | an image acquisition and processing software developed<br>by ZEISS | ZEISS ZEN | A user interface software integrated with Zeiss LSM900 microscope |
| Google<br>Chrome<br>interface | CATMAID:<br>collaborative<br>annotation toolkit for<br>massive amounts of<br>image data | [52] |  | Google Chrome<br>interface used for<br>TEM reconstruction<br>of neural circuits |
| Software | Fiji (Fiji Is Just ImageJ) | [53] |  | an open-source<br>image processing<br>package based on<br>ImageJ2 |
| Software | Adobe Illustrator | (Adobe, San<br>Jose, CA). |  | Was widely used for<br>generating figures |

### Muscle Calcium imaging, data quantification and analysis

A 25mm*25mm*2mm square pad made of 1.5% agarose gel was used as substrate of the larva during imaging. L2-L3 Larvae were washed in distilled water and individually placed onto the agarose gel pad, covered by a 22mm*22mm cover glass to gently constrain locomotion. The larvae were placed ventrolaterally to ensure VO muscles are always in the focal plane. All imaging was performed on a Zeiss LSM900 microscope with a 5X or a 10X objective. Up to 3 craw bouts from the same animal were used in quantification. For each wave muscles of two or three subsequent hemisegments were quantified. Calcium activity of individual muscles were quantified with custom MATLAB scripts. ROIs were put onto each muscle and adjusted on a frame-by-frame basis to track the muscle movement; overlapping regions and regions not clearly in focus were avoided. Since it was impossible to capture all 30 muscles in any hemisegment, we used segments containing at least one muscle representing each of the 4 co-active muscle groups defined by Zarin et. al. eLife 2019, and reused their PCA-based method to define the comprehensive activity peak, onset and offset of the entire segment. We aligned all crawl cycles to the onset and offset (corresponding to 30 and 70 on the 0-100 normalized time scale respectively) to exclude the effect of viable crawl speed. DF/F of each muscle was calculated as (F-F0)/F0, where F0 was defined as the tenth percentile value of the first 1/3 of the measurement. For boxplots, individual muscle activity onset was defined as the time point that muscle activity (DF/F) increased to 20% of the total DF/F increment (defined as the difference of peak value and minimum value before peak); onset points for normalized time and real time (second) were calculated independently. VO muscle onset delay (compared to the earliest muscle) is calculated as the difference between the onset time of individual VO muscles and the smallest muscle onset time within the same segment.

### Fly husbandry

All flies were reared in a 25°C room at 50% relative humidity with a 12 hr light/dark cycle. All comparisons between groups were based on studies with flies grown, handled, and tested together. For crosses involving UAS-GtACR1.d.EYFP, animals were reared at constant dark to avoid unwanted GtACR activation by ambient light. Larval offspring from these crosses were fed with all trans retinal (ATR) and were reared constant dark until they were ready to be imaged under the confocal microscope.

### PMN loss of function experiments

All fly stocks used for PMN loss of function experiments are listed in key resources table above. Briefly, to silence any given PMN (i.e., A06c, A27h, or A18b3), the Gal4 or split Gal4 targeting that PMN was crossed to 44H10::GCaMP6f, UAS-GtACR1.d.EYFP or 44H10::GCaMP6f, UAS-Kir2.1-EGFP. Offspring from these crosses were used for larval locomotion studies. For control groups, 44H10::GCaMP6f, UAS-GtACR1.d.EYFP or 44H10::GCaMP6f, UAS-Kir2.1-EGFP were corssed to w1118 strain and muscle imaging was done in their larval progeny.

Crosses and offspring involving UAS-Kir2.1-EGFP were reared in a 25°C room at 50% relative humidity with a 12 hr light/dark cycle. Crosses and offspring involving UAS-GtACR1.d.EYFP were wrapped by aluminum foil and were reared in a 25°C room at constant dark to avoid any unwanted exposure to ambient light. To fully sensitize GtACR1 to the confocal laser light, larval offspring were fed with all trans retinal (ATR) the night before the experiment day and were kept at dark until they were imaged by confocal. 100 mM All-trans-retinal (ATR) stock solution was made by dissolving 100 mg of all-trans-retinal powder (Sigma Aldrich R2500-100MG) into 3.52 ml 100% EtOH, and were divided into aliquots of 100 μl. Larval offspring were raised in *Drosophila* vials containing fly food. For each vial, 200 microliter of 0.5 mM ATR solution (1 microliter ATR stock solution dissolved in 199 microliter of H2O) was added the night before the optogenetic experiments.

### Calcium imaging in neurons

Dual-color neuronal imaging were done in freshly dissected brains. L2-L3 brains were dissected in HL3.1 saline [54] and mounted on 12mm round Poly-D-Lysine Coverslips and imaged on a Zeiss LSM900 confocal microscope. For dual-color imaging, two neurons differentially expressing GCaMP6m and jRCaMP1b were imaged in the same focal plane and ROIs were placed on individual neurons. Quantification of fluorescence activity was done on Zeiss Zen software and imported to MATLAB for analysis. Data from different fictive crawl cycles was aligned based on the activity of reference MNs (i.e,. CQ-LexA, LexAop-GCaMP6m). The 30% and 70% of each fictive crawl cycle was defined as the width (half height) of MN activity peak and the jRCaMP activity (PMN of the same segment) was aligned to GCaMP6m.

### Statistics

All statistical analysis, including t-tests and one-way ANOVA were performed with R 4.0.3 in RStudio (R Core Team, R Foundation for Statistical Computing, Vienna, Austria, and RStudio Team, RStudio, Inc., Boston, MA) and MATLAB R2021b (The MathWorks, Inc., Natick, MA). Numerical data in graphs show measurements of individual muscles (scatter dots on boxplots in Figures 1D, 4C-D, 6C-D, means (midline of boxplots in Figures 1D, 4C-D, 6C-D or means ± SD (Figures 1D, 3A-C, 4C-D, 5B, 6C-D. The number of replicates (n, representing numbers of animals, individual muscles, or crawl bouts) in indicated for each data set in the corresponding legend. Sample size was pre-determined by a power analysis with effect size d = 0.5, α = 0.05 and power = 0.8 in the freeware G*Power3.1 (Faul et al. 2009) [51].

### Electron microscopy and CATMAID reconstructions

The TEM volume is for a newly hatched first instar (L1) larval nervous system (Ohyama et al., 2015). This dataset is available by request from Albert Cardona (Cambridge University). Neurons were reconstructed in CATMAID (Saalfeld et al., 2009) [52] using a Google Chrome browser as previously described (Ohyama et al., 2015). Figure 2 was generated using CATMAID 3D widget, Python script, and Adobe Illustrator (Adobe, San Jose, CA).

## Data availability

The source data for figures 1-6 as well as customized MATLAB scripts for data processing, analysis and plotting are included with this submission. All data, including raw confocal imaging data of larval muscles and isolated brains, quantification of muscle or neuron activity, and any other data that generated or analyzed during this study, will be included with this manuscript once it is accepted for publication.

## Acknowledgements

We thank Jim Truman for unpublished fly lines targeting A18b3 and A06c neurons. We thank Chris Q Doe, Wanhe Li, and Lewis Sherer for comments on early versions of the manuscript. We thank Chris Q Doe, Todd Laverty, Gerry Rubin for fly stocks, We thank Albert Cardona, Richard D. Fetter, and the HHMI Janelia Fly EM Project Team for providing the raw data of the whole CNS EM volume. We thank Akira Fushiki, Eri Hasegawa, and Maarten Zwart for annotating neurons, and Keiko Hirono for generating transgenic constructs. AAZ and YH were supported by institutional startup fund provided to AAZ by Texas A&M University.

---

**Figure 5 figure supplement 1.**
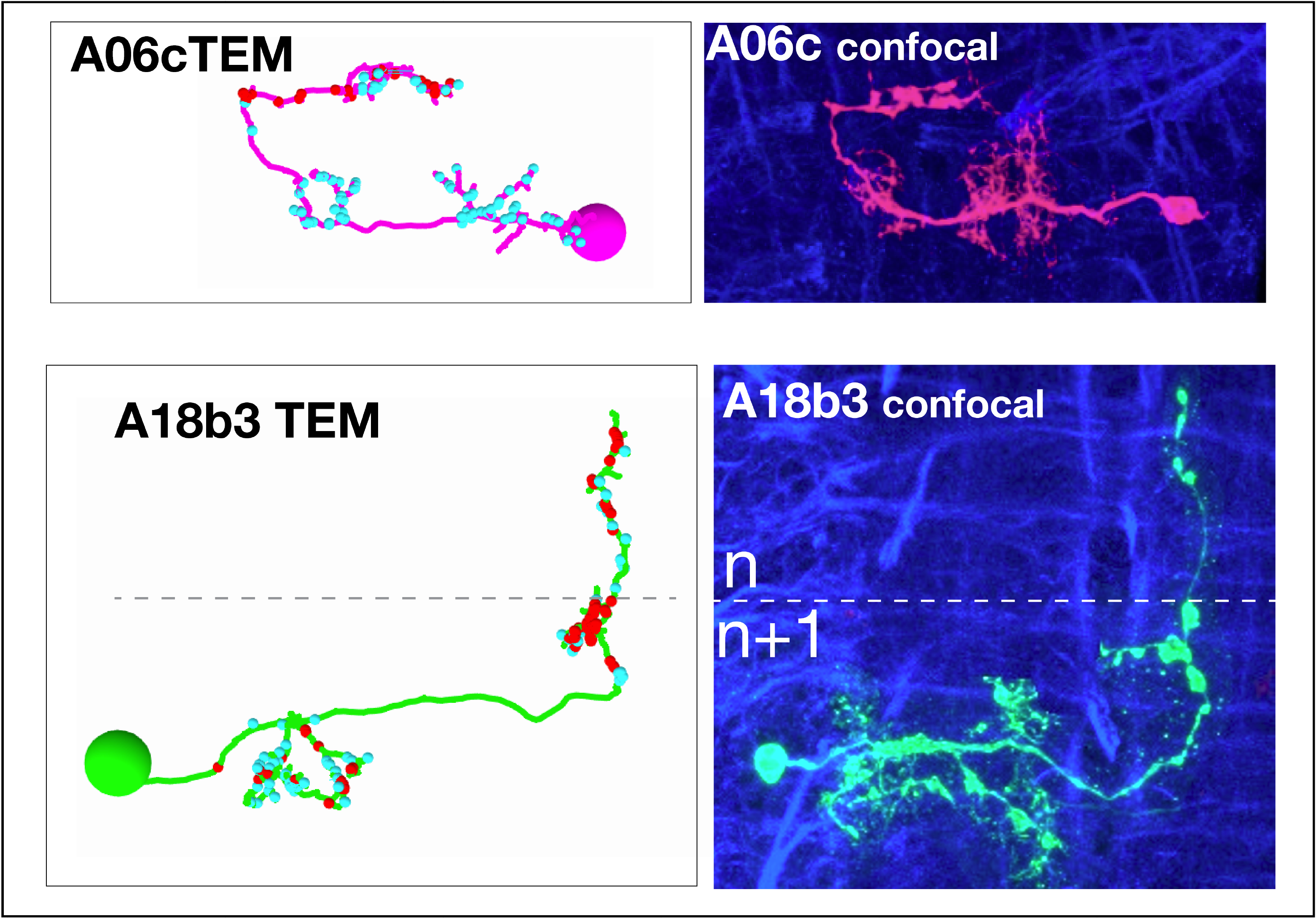
A06c and A18b3 PMN morphology by light and electron microscopy (TEM). Top: Dorsal view of individual A06c and A18b3 PMNs in a second instar larval CNS by light microscopy (R94E10 > MCFO). Bottom: of individual A06c and A18b3 PMNs in a first instar larva in the TEM reconstruction. Cyan dots, post-synaptic sites; red dots, pre-synaptic sites. Anterior, left. Midline, dashed line.

**Video 1. The VO muscles are activated earlier in backward than in forward locomotion**. Muscle calcium imaging in an intact wild-type animal performing forward and then backward locomotion. White arrowheads point at VO muscles as they are being activated in each segment. Anterior is left and dorsal is up.

**Video 2. A06c PMN has different firing patterns in forward and backward locomotion**. Dual color calcium imaging in an isolated brain performing fictive backward and then forward locomotion. Top panel shows the reference MNs expressing GCaMP6m and bottom panel shows A06c PMNs expressing RCaMP1b. White arrowheads point at MNs and A06c PMN as they are actively firing in a single segment. Anterior is left. During backward locomotion (from 0:00 to 0:13),, A06c PMN fires only once after MNs in its corresponding segment. During forward locomotion (from 0:13 to 0:31), A06c PMN fires twice, once before and once after MNs in its corresponding segment.

**Video 3. A27h PMN is active in forward but not backward locomotion**. Dual color calcium imaging in an isolated brain performing fictive backward and then forward locomotion. Top panel shows the reference MNs expressing GCaMP6m and bottom panel shows A27h PMNs expressing RCaMP1b. White arrowheads point at MNs and A27h PMN as they are actively firing in a single segment. Anterior is left. During forward locomotion (from 0:00 to 0:13), A27h PMN fires almost synchronously with MNs in its corresponding segment. During backward locomotion (from 0:13 to 0:23), A27h PMN has no activity. In addition to A27h, 36G02-Gal4 line hits another neuron known as A03g and M neuron [46, 55]. The axons shown in red channel belong only to A27h, as the areas containing A03g/M neuron were cropped out. The data from this animal has been previously published in another paper published by the same author [12].

**Video 4. Loss of function (LOF) of A06c PMN leads to premature VO muscle activation in forward locomotion**. Muscle calcium imaging in an A06c silenced animal performing forward locomotion. White arrowheads point at VO muscles as they are being activated in each segment. Anterior is left and dorsal is up.

**Video 5. Loss of function (LOF) of A27h PMN leads to premature VO muscle activation in forward locomotion**. Muscle calcium imaging in an A27h silenced animal performing forward locomotion. White arrowheads point at VO muscles as they are being activated in each segment. Anterior is left and dorsal is up.

**Video 6. A18b3 PMN is active in forward but not backward locomotion**. Dual color calcium imaging in an isolated brain performing fictive backward and then forward locomotion. Top panel shows the reference MNs expressing GCaMP6m and bottom panel shows A18b3 PMNs expressing RCaMP1b. White arrowheads point at MNs and A18b3 PMN as they are actively firing in a single segment. Anterior is left. During forward locomotion (from 0:00 to 0:12), A18b3 PMN fires simultaneously with MNs in its corresponding segment. During backward locomotion (from 0:12 to 0:26), A18b3 PMN has no activity.

**Video 7. Loss of function (LOF) of A18b3 PMN leads to premature VO muscle activation in forward locomotion**. Muscle calcium imaging in an A18b3 silenced animal performing forward locomotion. White arrowheads point at VO muscles as they are being activated in each segment. Anterior is left and dorsal is up.

**Figure 1—source data 1**

**Figure 2—source data 1**

**Figure 3—source data 1**

**Figure 4—source data 1**

**Figure 5—source data 1**

**Figure 6—source data 1**

**Figure 7—source data 1**

## References

1. Talpalar, A.E., et al., Dual-mode operation of neuronal networks involved in left-right alternation. Nature, 2013. 500(7460): p. 85–8.

2. Bellardita, C. and O. Kiehn, Phenotypic characterization of speed-associated gait changes in mice reveals modular organization of locomotor networks. Curr Biol, 2015. 25(11): p. 1426–36.

3. Liao, J.C. and J.R. Fetcho, Shared versus specialized glycinergic spinal interneurons in axial motor circuits of larval zebrafish. J Neurosci, 2008. 28(48): p. 12982–92.

4. Green, C.S. and S.R. Soffe, Transitions between two different motor patterns in Xenopus embryos. J Comp Physiol A, 1996. 178(2): p. 279–91.

5. Berkowitz, A., A. Roberts, and S.R. Soffe, Roles for multifunctional and specialized spinal interneurons during motor pattern generation in tadpoles, zebrafish larvae, and turtles. Front Behav Neurosci, 2010. 4: p. 36.

6. Hooper, J.E., Homeotic gene function in the muscles of Drosophila larvae. Embo j, 1986. 5(9): p. 2321–2329.

7. Bate, M., The embryonic development of larval muscles in Drosophila. Development, 1990. 110(3): p. 791–804.

8. Bate, M. and E. Rushton, Myogenesis and muscle patterning in Drosophila. C R Acad Sci III, 1993. 316(9): p. 1047–61.

9. Landgraf, M., et al., The origin, location, and projections of the embryonic abdominal motorneurons of Drosophila. J Neurosci, 1997. 17(24): p. 9642–55.

10. Zwart, M.F., et al., Selective Inhibition Mediates the Sequential Recruitment of Motor Pools. Neuron, 2016. 91(3): p. 615–28.

11. Zarin, A.A. and J.P. Labrador, Motor axon guidance in Drosophila. Semin Cell Dev Biol, 2017.

12. Zarin, A.A., et al., A multilayer circuit architecture for the generation of distinct locomotor behaviors in Drosophila. Elife, 2019. 8.

13. Ohyama, T., et al., A multilevel multimodal circuit enhances action selection in Drosophila. Nature, 2015. 520(7549): p. 633–9.

14. Jovanic, T., et al., Competitive Disinhibition Mediates Behavioral Choice and Sequences in Drosophila. Cell, 2016. 167(3): p. 858–870 e19.

15. Burgos, A., et al., Nociceptive interneurons control modular motor pathways to promote escape behavior in Drosophila. Elife, 2018. 7.

16. Carreira-Rosario, A., et al., MDN brain descending neurons coordinately activate backward and inhibit forward locomotion. Elife, 2018. 7.

17. Clark, M.Q., et al., Neural circuits driving larval locomotion in Drosophila. Neural Dev, 2018. 13(1): p. 6.

18. Fushiki, A., et al., A circuit mechanism for the propagation of waves of muscle contraction in Drosophila. Elife, 2016. 5.

19. Lnenicka, G.A. and H. Keshishian, Identified motor terminals in Drosophila larvae show distinct differences in morphology and physiology. J Neurobiol, 2000. 43(2): p. 186–97.

20. Lu, Z., et al., High-Probability Neurotransmitter Release Sites Represent an Energy-Efficient Design. Curr Biol, 2016. 26(19): p. 2562–2571.

21. Newman, Z.L., et al., Input-Specific Plasticity and Homeostasis at the Drosophila Larval Neuromuscular Junction. Neuron, 2017. 93(6): p. 1388-1404.e10.

22. Genc, O. and G.W. Davis, Target-wide Induction and Synapse Type-Specific Robustness of Presynaptic Homeostasis. Curr Biol, 2019. 29(22): p. 3863–3873 e2.

23. Perez-Moreno, J.J. and C.J. O’Kane, GAL4 Drivers Specific for Type Ib and Type Is Motor Neurons in Drosophila. G3 (Bethesda), 2019. 9(2): p. 453–462.

24. Aponte-Santiago, N.A. and J.T. Littleton, Synaptic Properties and Plasticity Mechanisms of Invertebrate Tonic and Phasic Neurons. Front Physiol, 2020. 11: p. 611982.

25. Aponte-Santiago, N.A., et al., Synaptic Plasticity Induced by Differential Manipulation of Tonic and Phasic Motoneurons in Drosophila. J Neurosci, 2020. 40(33): p. 6270–6288.

26. Wang, Y., et al., Structural and Functional Synaptic Plasticity Induced by Convergent Synapse Loss in the Drosophila Neuromuscular Circuit. J Neurosci, 2021. 41(7): p. 1401–1417.

27. Mohammad, F., et al., Optogenetic inhibition of behavior with anion channelrhodopsins. Nat Methods, 2017. 14(3): p. 271–274.

28. Hasegawa, E., J.W. Truman, and A. Nose, Identification of excitatory premotor interneurons which regulate local muscle contraction during Drosophila larval locomotion. Sci Rep, 2016. 6: p. 30806.

29. Kuo, D.H., et al., A tale of two leeches: Toward the understanding of the evolution and development of behavioral neural circuits. Evol Dev, 2020. 22(6): p. 471–493.

30. Popescu, I.R. and W.N. Frost, Highly dissimilar behaviors mediated by a multifunctional network in the marine mollusk Tritonia diomedea. J Neurosci, 2002. 22(5): p. 1985–93.

31. Butt, S.J., R.M. Harris-Warrick, and O. Kiehn, Firing properties of identified interneuron populations in the mammalian hindlimb central pattern generator. J Neurosci, 2002. 22(22): p. 9961–71.

32. Gosgnach, S., et al., Delineating the Diversity of Spinal Interneurons in Locomotor Circuits. J Neurosci, 2017. 37(45): p. 10835–10841.

33. Ampatzis, K., et al., Separate microcircuit modules of distinct v2a interneurons and motoneurons control the speed of locomotion. Neuron, 2014. 83(4): p. 934–43.

34. Menelaou, E. and D.L. McLean, Hierarchical control of locomotion by distinct types of spinal V2a interneurons in zebrafish. Nat Commun, 2019. 10(1): p. 4197.

35. Rybak, I.A., K.J. Dougherty, and N.A. Shevtsova, Organization of the Mammalian Locomotor CPG: Review of Computational Model and Circuit Architectures Based on Genetically Identified Spinal Interneurons(1,2,3). eNeuro, 2015. 2(5).

36. Ziskind-Conhaim, L. and S. Hochman, Diversity of molecularly defined spinal interneurons engaged in mammalian locomotor pattern generation. J Neurophysiol, 2017. 118(6): p. 2956–2974.

37. Crone, S.A., et al., In mice lacking V2a interneurons, gait depends on speed of locomotion. J Neurosci, 2009. 29(21): p. 7098–109.

38. Dougherty, K.J., et al., Locomotor rhythm generation linked to the output of spinal shox2 excitatory interneurons. Neuron, 2013. 80(4): p. 920–33.

39. Milner-Brown, H.S., R.B. Stein, and R. Yemm, The orderly recruitment of human motor units during voluntary isometric contractions. J Physiol, 1973. 230(2): p. 359–70.

40. Henneman, E., The size-principle: a deterministic output emerges from a set of probabilistic connections. J Exp Biol, 1985. 115: p. 105–12.

41. Gabriel, J.P., et al., Control of flexor motoneuron activity during single leg walking of the stick insect on an electronically controlled treadwheel. J Neurobiol, 2003. 56(3): p. 237–51.

42. Mendell, L.M., The size principle: a rule describing the recruitment of motoneurons. J Neurophysiol, 2005. 93(6): p. 3024–6.

43. Hill, A.A. and D. Cattaert, Recruitment in a heterogeneous population of motor neurons that innervates the depressor muscle of the crayfish walking leg muscle. J Exp Biol, 2008. 211(Pt 4): p. 613–29.

44. Azevedo, A.W., et al., A size principle for recruitment of Drosophila leg motor neurons. Elife, 2020. 9.

45. Schneider-Mizell, C.M., et al., Quantitative neuroanatomy for connectomics in Drosophila. Elife, 2016. 5.

46. Kohsaka, H., et al., Regulation of forward and backward locomotion through intersegmental feedback circuits in Drosophila larvae. Nat Commun, 2019. 10(1): p. 2654.

47. Takagi, S., et al., Divergent Connectivity of Homologous Command-like Neurons Mediates Segment-Specific Touch Responses in Drosophila. Neuron, 2017. 96(6): p. 1373-1387.e6.

48. Li, H.H., et al., A GAL4 driver resource for developmental and behavioral studies on the larval CNS of Drosophila. Cell Rep, 2014. 8(3): p. 897–908.

49. Dionne, H., et al., Genetic Reagents for Making Split-GAL4 Lines in Drosophila. Genetics, 2018. 209(1): p. 31–35.

50. Giachello, C.N.G., et al., Electrophysiological validation of monosynaptic connectivity between premotor interneurons and the aCC motoneuron in the Drosophila larval CNS. J Neurosci, 2022.

51. Faul, F., et al., Statistical power analyses using G*Power 3.1: tests for correlation and regression analyses. Behav Res Methods, 2009. 41(4): p. 1149–60.

52. Saalfeld, S., et al., CATMAID: collaborative annotation toolkit for massive amounts of image data. Bioinformatics, 2009. 25(15): p. 1984–6.

53. Schindelin, J., et al., Fiji: an open-source platform for biological-image analysis. Nat Methods, 2012. 9(7): p. 676–82.

54. Feng, Y., A. Ueda, and C.F. Wu, A modified minimal hemolymph-like solution, HL3.1, for physiological recordings at the neuromuscular junctions of normal and mutant Drosophila larvae. J Neurogenet, 2004. 18(2): p. 377–402.

55. Zeng, X., et al., An electrically coupled pioneer circuit enables motor development via proprioceptive feedback in Drosophila embryos. Curr Biol, 2021. 31(23): p. 5327–5340 e5.

